# Switching secretory pathway direction for organelle acquisition in plants

**DOI:** 10.1101/2020.03.02.956961

**Authors:** Takehiko Kanazawa, Hatsune Morinaka, Kazuo Ebine, Takashi L. Shimada, Sakiko Ishida, Naoki Minamino, Katsushi Yamaguchi, Shuji Shigenobu, Takayuki Kohchi, Akihiko Nakano, Takashi Ueda

## Abstract

Eukaryotic cells acquired novel organelles during evolution through mechanisms that remain largely obscure. The existence of the unique oil body compartment is a synapomorphy of liverworts that represents lineage-specific acquisition of this organelle during evolution, although its origin, biogenesis, and physiological function are yet unknown. We found that two Syntaxin 1 paralogs in the liverwort, *Marchantia polymorpha*, are distinctly targeted to forming cell plates and the oil body, suggesting these structures share some developmental similarity. Oil body formation is under the regulation of an ERF/AP2-type transcription factor and loss of the oil body increased *M. polymorpha* herbivory. These findings highlight a common strategy for the acquisition of organelles with distinct functions in plants, via periodical switching in secretion direction depending on cellular phase transition.

## Main Text

Eukaryotic cells originated from prokaryotes by expanding their endomembrane network during evolution, with the last eukaryotic common ancestor (LECA) likely possessing a complex set of organelles ^1^. New membrane trafficking pathways were added to the LECA endomembrane network, some of which were secondarily lost in a lineage-specific manner, resulting in divergent and organism-specific membrane trafficking systems and organelle compositions of extant eukaryotes ^2-4^. However, it remains mostly unknown how organelles were acquired during evolution. Also in the plant lineage, several organelles and organelle functions have been uniquely acquired during evolution. For example, the plant vacuole harbours a unique function that is not shared with the animal lysosome and the yeast vacuole: storage of proteins ^5^. This vacuolar function is fulfilled through the plant-unique vacuolar trafficking system, which comprises multiple vacuolar transport pathways involving plant-unique machinery components acquired during plant evolution ^6^. The cell plate, which is formed during the mitotic phase to accomplish cytokinesis in land plants, is also a prominent example of plant-specific cellular structures/organelles ^7^.

One of the basal-most land plant lineages, liverworts, also possess a unique organelle, the oil body, existence of which is a synapomorphy of this lineage. The liverwort oil body contains bioactive compounds such as sesquiterpenoids and cyclic bisbibenzyl compounds and is not related to the oil body that stores neutral lipids in storage organs like seeds and fruits (i.e. lipid body or oleosome). About 90% of liverwort species have this organelle; however, its origin, biogenesis, and physiological function remain unclear with controversial origins proposed from microscopic observations ^8-11^, although the first description of the liverwort oil body dates back to 1834 ^12^. Through the systematic analysis of SNARE proteins in the liverwort, *Marchantia polymorpha* (hereafter referred to as Marchantia), we identified an oil body-resident protein, MpSYP12B ^13^, which is a homolog of animal Syntaxin 1 that acts in the final step of exocytosis. Functions of SYP1 members to which MpSYP12B belongs dramatically diversified during plant evolution, suggesting expanded secretory trafficking systems in plants. Arabidopsis SYP1 consists of three subgroups, SYP11, SYP12, and SYP13. The SYP13 group mediates constitutive secretion, whereas the other groups are involved in plant-specific higher ordered functions; for example, KNOLLE (also known as SYP111) is specifically involved in membrane fusion at forming cell plates, and PEN1 (also known as SYP121) is required for intact penetration resistance against powdery mildew fungi and regulation of the ion channel activity ^14-19^. For more insight into the functional diversification of SYP1 members in Marchantia and the mechanism of oil body biogenesis, we further characterized four SYP1 members in this organism ^13,20^.

### MpSYP12A is important for cell plate formation

Our phylogenetic analysis suggested that MpSYP1 members belong to two distinct subgroups: the SYP13 group (MpSYP13A and 13B) and the SYP11/12 group (MpSYP12A and 12B) that contains SYP11 and SYP12 groups in seed plants ^21^ (Fig. S1A). MpSYP12A, 13A and 13B were ubiquitously expressed and localized to the plasma membrane (PM) in thallus tissues, whereas MpSYP12B exhibited specific expression in a subpopulation of thallus cells (Fig S1B-E). Using some ubiquitously expressed mCitrine-MpSYP1 protein fusion constructs driven by their own regulatory elements, we detected specific localization of MpSYP12A at forming cell plates associated with the phragmoplast that rapidly stained with FM4-64 (Figs. 1A-F, S1F, and S1G). This localization suggested that MpSYP12A could be a functional counterpart of KNOLLE in Arabidopsis, which was further verified genetically.

**Fig. 1.**
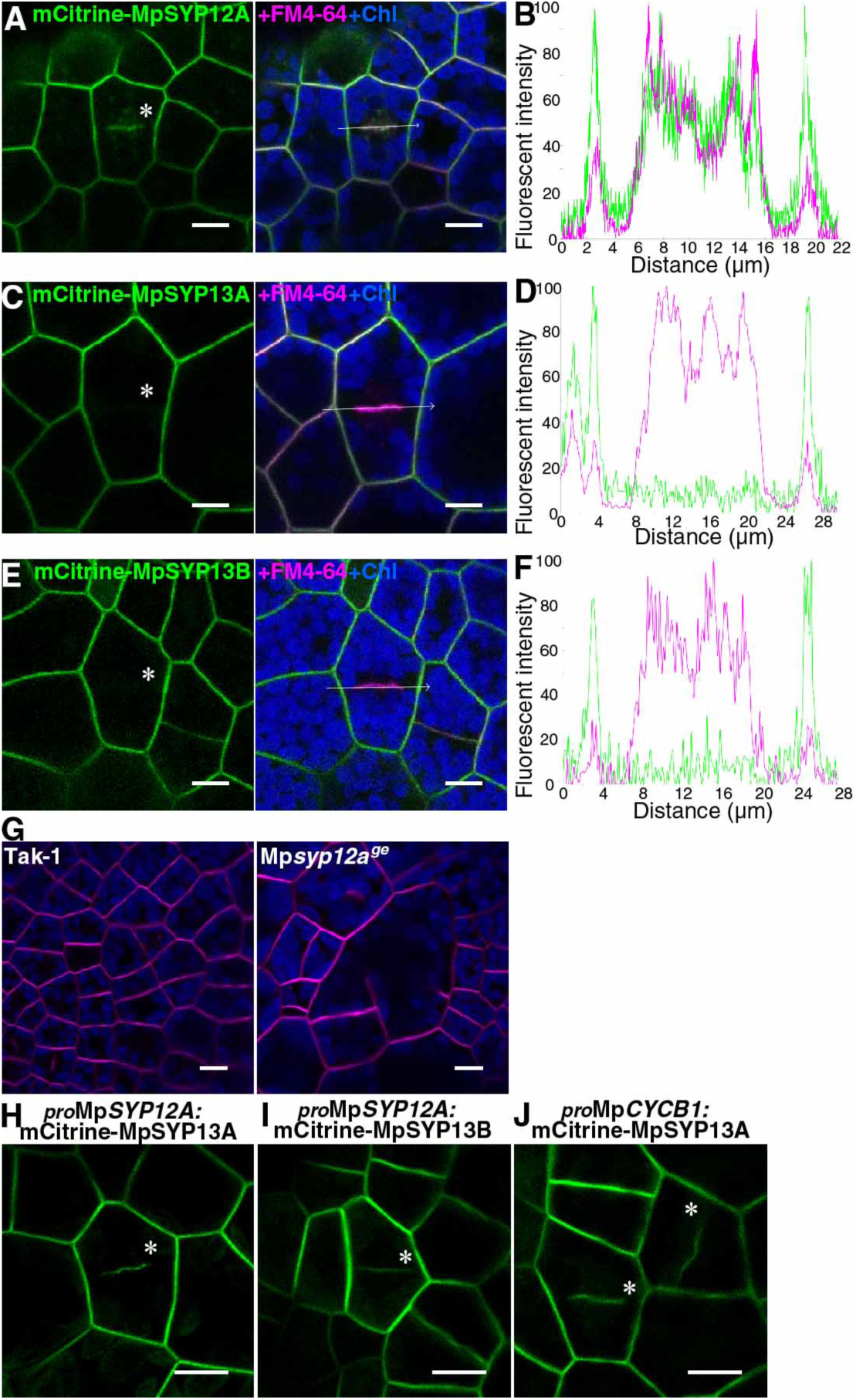
MpSYP12A is important for cell plate formation in *M. polymorpha* thallus cells. (A, C, and E) Marchantia thallus cells expressing mCitrine-MpSYP12A (A), mCitrine-MpSYP13A (C), and mCitrine-MpSYP13B (E) stained with FM4-64. (B, D, and F) Line graphs showing relative fluorescence intensities from mCitrine and FM4-64 along the white arrows in panels A (B), C (D), and E (F). (G) Thallus cells of Tak-1 (left) and a putative knock-out mutant (right) stained with FM4-64. (H and I) Dividing thallus cells expressing mCitrine-MpSYP13A (H) or mCitrine-MpSYP13B (I) driven by the Mp*SYP12A* promoter. (J) Dividing thallus cells expressing mCitrine-MpSYP13A under the regulation of the Mp*CYCB1* promoter. Asterisks indicate forming cell plates. Green, magenta, and blue pseudo-colors indicate fluorescence from mCitrine, FM4-64, and chlorophyll, respectively. Bars = 10 µm.

We tried to generate a complete Mp*syp12a* knockout mutant by genome editing (Fig. S1H) but were not successful, probably because it is an essential gene. However, we succeeded in isolating chimeric plants comprising mutated and wild-type cells, in which we frequently observed enlarged cells with cell wall stabs, suggesting that MpSYP12A is required for cell plate formation during cytokinesis similar to the function of KNOLLE (Fig. 1G). The *KNOLLE* promoter is sufficient to target non-cytokinesis-specific SYP132 to forming cell plates ^22^. Similarly, mCitrine-tagged MpSYP13A and 13B were localized to cell plates and the PM when expressed under the Mp*SYP12A* promoter (Fig. 1H, I). Furthermore, a similar localization was also observed when mCitrine-MpSYP13 was expressed by the Mp*CYCB1* promoter (Figs. 1J and S1I), indicating that the cell cycle-dependent transcriptional regulation is also critical for cell plate targeting of SYP1 proteins in Marchantia. We also found that clathrin light chain (CLC) tagged with mCitrine localized to the PM and *trans*-Golgi network in non-dividing thallus cells, as well as on forming cell plates in Marchantia as has been reported in other plants (Fig. S1J, K) ^23-25^. These results strongly suggested that fundamental mechanisms of cell plate formation were conserved during land plant evolution.

### The oil body membrane shares common properties with the plasma membrane

Distinct from the other MpSYP1 members, MpSYP12B exhibits specific expression in oil body cells and localizes to the oil body membrane ^13^. Oil body cells expressing a _*pro*_Mp*SYP12B:*2×YFP construct indicated that MpSYP12B distributed around meristematic regions in young thalli, which was observed using light-sheet microscopy (Fig. 2A; Supplementary movie S1). These results suggest that oil body formation occurs around meristematic regions and into thalli during the growth of thallus tissues. mCitrine-MpSYP12B expressed under its own regulatory elements localized to the oil body membrane with a faint signal on the PM (Fig. 2B), which was also confirmed by immuno-EM analysis using an anti-GFP antibody (Fig. 2C, D). We then tested whether organelle markers for the endoplasmic reticulum (MpSEC20, MpUSE1A, MpSEC22, mRFP-HDEL, and GFP-HDEL), Golgi apparatus (MpGOS11 and MpSFT1), *trans*-Golgi network (MpSYP6A and MpSYP4), and tonoplast (MpSYP2 and MpVAMP71) were targeted to the oil body membrane, none of which were detected (Figs. 2E and S2B, C). Using transmission electron microscopy, we found that clathrin-coated vesicles formed from the oil body membrane, and emergence and disappearance of clathrin-positive foci at the oil body membrane was also observed in transgenic plants expressing Citrine-tagged MpCLC (Fig. 2G; Supplementary movie S2).

**Fig. 2.**
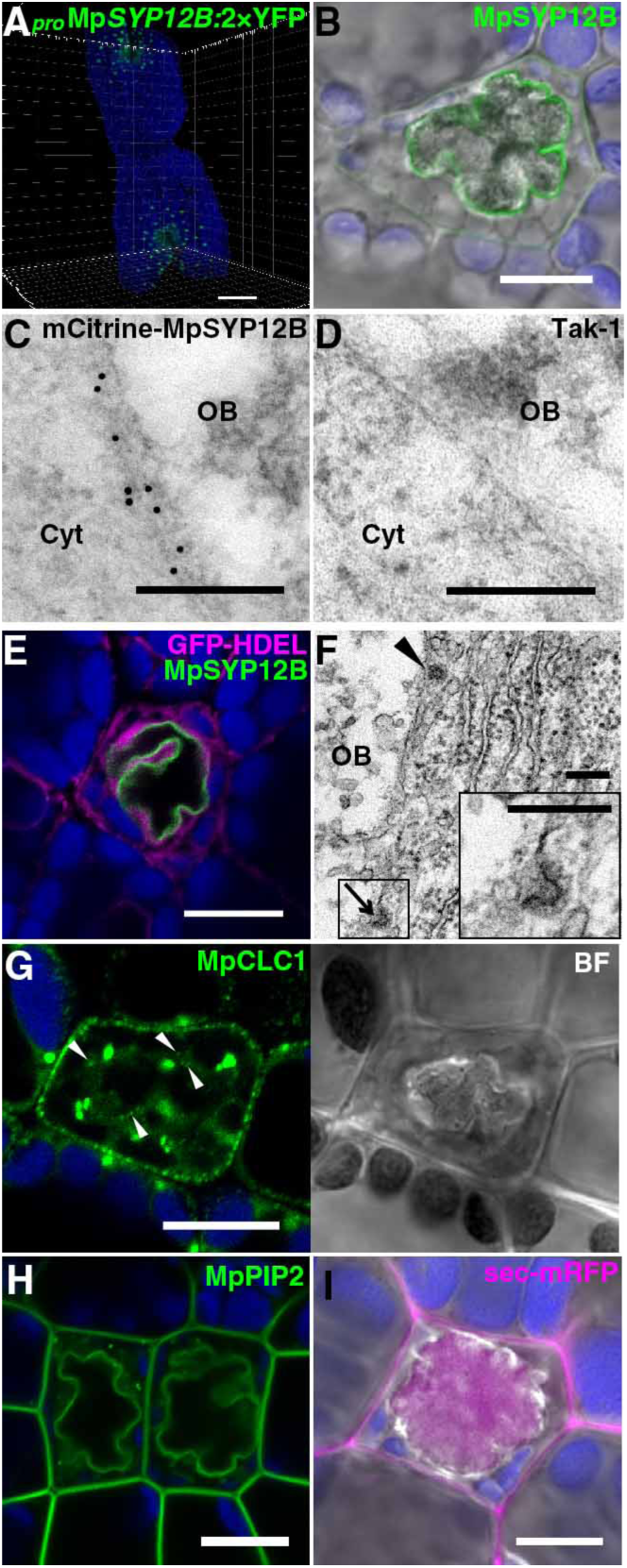
The oil body membrane shares common properties with the plasma membrane. (A) A five-day-old thallus expressing 2×Citrine (YFP) under the regulation of the Mp*SYP12B* promoter observed with a light sheet microscope. (B) An oil body cell expressing mCitrine-MpSYP12B under its own regulatory elements. The bright field image was merged. (C and D) Immunoelectron micrographs of oil body cells in thalli of Tak-1 with (C) or without (D) mCitrine-MpSYP12B expression. Cyt, cytosol, and OB, oil body. (E) An oil body cell expressing GFP-HDEL (magenta) and mCitrine-MpSYP12B (green). (F) An electron micrograph of the oil body cell. The arrowhead and arrow indicate a clathrin-coated vesicle and a clathrin-coated pit, respectively. The magnified image of the boxed region is shown in the inset. (G) An oil body cell expressing MpCLC1-Citrine. Arrowheads indicate clathrin-coated vesicles/pits on the oil body membrane. BF, bright field image. (H) Oil body cells expressing mCitrine-MpPIP2 under its own regulatory elements. (I) An oil body cell expressing sec-mRFP driven by the Mp*EF1α* promoter. The bright field image was merged. The blue pseudo-color indicates chlorophyll autofluorescence. Bars = 500 µm in (A), 10 µm in (B), (E), and (G - I), and 200 nm in (C), (D), and (F).

Given that clathrin-mediated endocytosis occurs at the PM and SYP1 members generally function on the PM, the oil body membrane could share similar characteristics with the PM. This hypothesis was supported by dual localization of the PM proteins, PM-type aquaporin MpPIP2 (Fig. 2H) and PM-resident SNARE MpSYP13A (Fig. 3A) driven by their own regulatory elements. The plasma membrane-like nature of the oil body membrane and the existence of the SYP1 member that is homologous to KNOLLE suggested that the oil body could be formed by the fusion of secretory vesicles similar to the cell plate. Thus, the luminal space of the oil body should be topologically equivalent to the extracellular space, which was tested by expressing a general secretion marker sec-mRFP, which is composed of the signal peptide for ER translocation and mRFP under the constitutive Mp*EF1α* promoter. In non-oil body cells, mRFP fluorescence was detected only in the extracellular space (Fig. S2D). However, mRFP accumulated in the oil body in addition to the extracellular space in oil body cells, demonstrating equivalent topology between the lumen of the oil body and extracellular space (Fig. 2I). Unlike MpSYP12A and KNOLLE, MpSYP12B loss of function did not result in a detectable abnormality in oil body formation and transport of MpSYP13A or sec-mRFP (Fig. S3), probably reflecting a functional redundancy between MpSYP12B and 13A similar to the partial functional redundancy of KNOLLE and SYP132 ^26^.

**Fig. 3.**
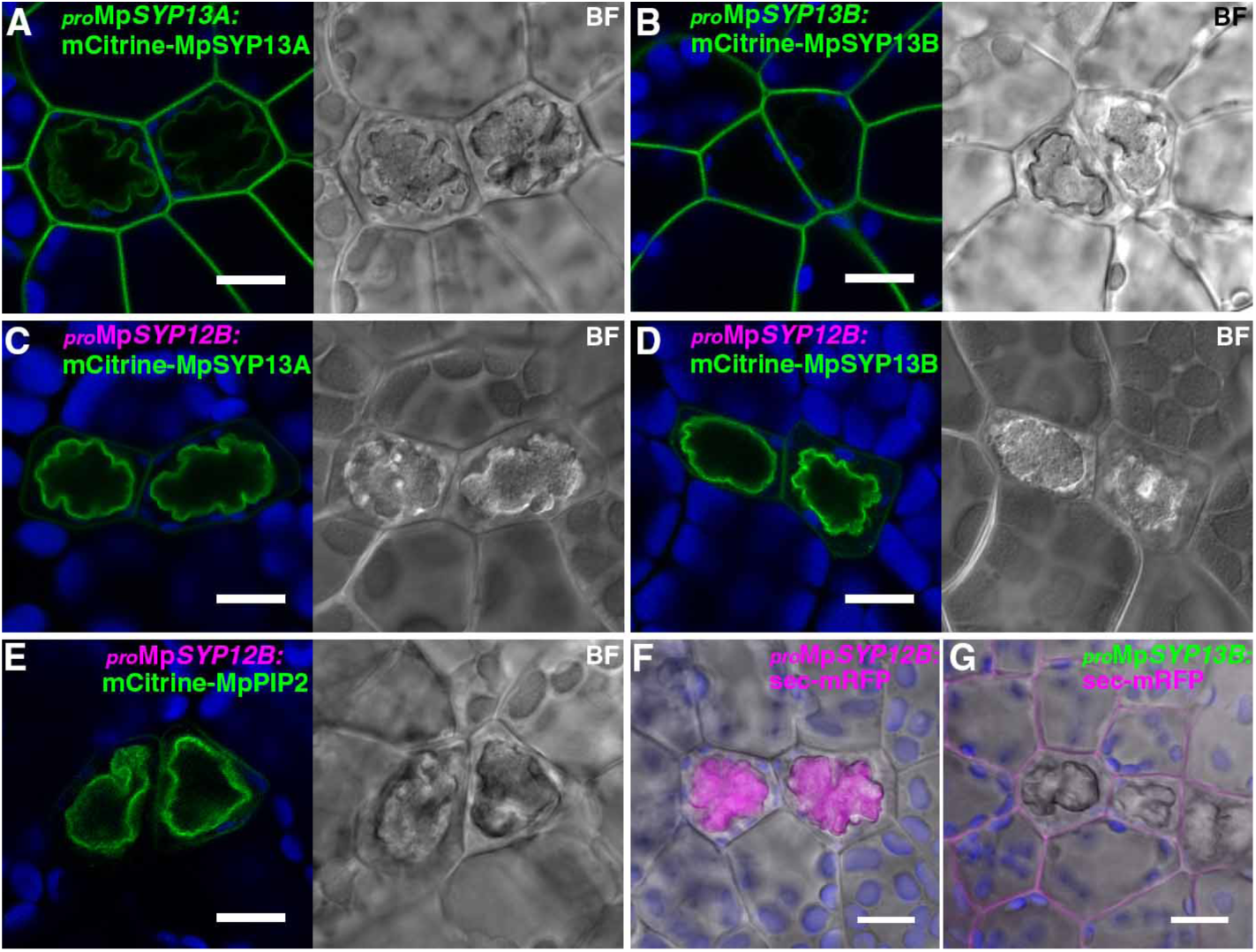
The oil body is formed by directional switching of the secretory pathway. (A and B) Thallus cells including oil body cells expressing mCitrine-MpSYP13A (A) or mCitrine- MpSYP13B (B) under their own regulatory elements. (C - E) Thallus cells including oil body cells expressing mCitrine-MpSYP13A (C), mCitrine-MpSYP13B (D), or mCitrine-MpPIP2 (E) under the Mp*SYP12B* promoter. BF, bright field images. (F and G) Thallus cells including oil body cells expressing sec-mRFP under the Mp*SYP12B* promoter (F) or the Mp*SYP13B* promoter (G). Bright field images are merged. Bars = 10 µm.

### The oil body is formed by directional switching of the secretory pathway

Distinct from MpSYP13A with the dual localization to the plasma and oil body membranes, the close homolog MpSYP13B was only localized to the PM in oil body cells (Fig. 3A, B). Surprisingly, both of these proteins were targeted predominantly to the oil body membrane when expressed under the regulation of the Mp*SYP12B* promoter (Fig. 3C, D). This effect was not restricted to SYP1 homologs; the PM-resident protein MpPIP2 and a general secretion cargo sec-mRFP were also targeted almost exclusively to the oil body when driven by the Mp*SYP12B* promoter (Fig. 3E, F). These results indicated that the secretory pathway is redirected inward and vesicles fuse with each other to form the oil body during the phase in which the Mp*SYP12B* promoter is active. In contrast, when Mp*SYP13B* promoter is active, the secretory pathway is directed to the PM and extracellular space, which was further confirmed by accumulation of sec-mRFP predominantly in the extracellular space when the construct was driven by the Mp*SYP13B* promoter (Fig. 3G). Other organelle markers did not change their localization even when expressed under the Mp*SYP12B* promoter (Fig. S2B, C). These results indicated that the switching of directions of the secretory pathway in oil body cells is regulated at the transcription level. The switching of secretory directions should take place periodically and repeatedly through oil body cell development, because we never observed oil body membrane localization for MpSYP13B at any developmental stages of oil body cells (Fig. 4A-C), and increase in the size of the oil body cell along with the increase in the oil body size was observed (Fig. 4D). Based on these findings, we propose the “oil body cycle hypothesis”, which states that Marchantia oil body cells cycle between two distinct cellular phases: the “PM phase” in which the secretory pathway is oriented to the PM and extracellular space, and the “oil body phase” when the secretory pathway is oriented to the oil body, with the phase transition under the regulation of a transcriptional regulatory system (Fig. 4E, F).

**Fig. 4.**
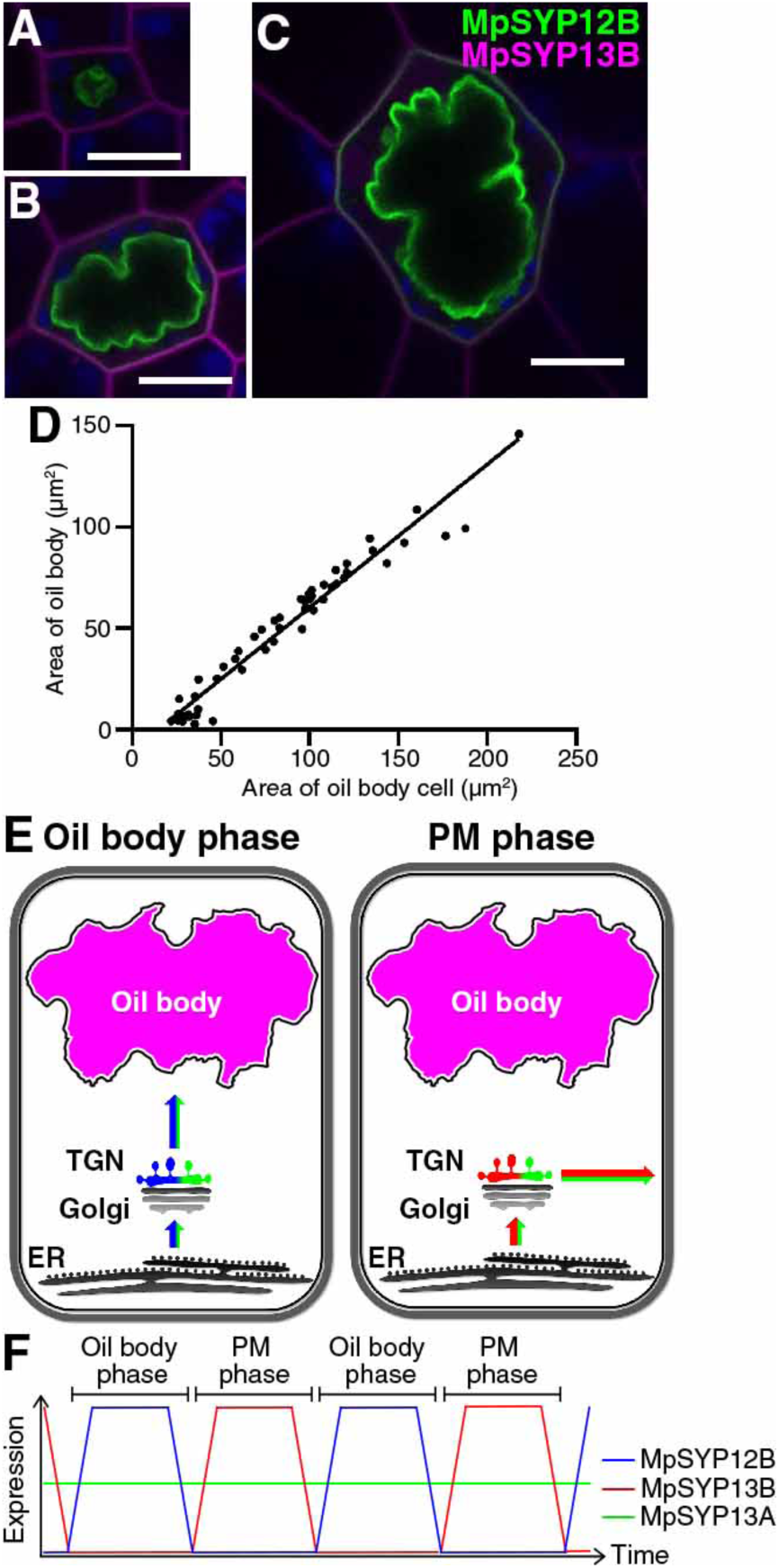
The oil body cycle hypothesis. (A - C) Oil body cells at distinct developmental stages expressing mCitrine-MpSYP12B and mRFP-MpSYP13B. Green, magenta, and blue pseudo-colors indicate mCitrine, mRFP, and autofluorescence from chlorophyll, respectively. Bars = 10 µm. (D) A scatter diagram of the areas of oil body cells on the X axis and the areas of oil bodies on the Y axis. R^2^ = 0.9547. (E and F) Schemes of the oil body cycle hypothesis. Newly synthesized PM proteins and secreted cargos are targeted to the oil body membrane in the oil body phase, and then to the PM and extracellular space during the PM phase (E). Expression levels of MpSYP12B and MpSYP13B periodically oscillate depending on the cell phases; whereas, MpSYP13A is constitutively expressed, resulting in dual localization at the oil body membrane and the PM (F).

### MpERF13 transcription factor regulates oil body formation

To investigate the regulatory system of oil body formation and oil body cycle, we screened mutants defective in oil body formation (Fig. S4A). From 48,825 T-DNA insertion lines, we identified a mutant (Mp*erf13*^*GOF*^) with an increased number of oil bodies. The mutant gemmae contained 334.4 ± 85.6 oil bodies (mean ± SD), whereas wild-type gemmae possessed 51.9 ± 8.7 oil bodies under our experimental conditions (Fig. 5A-E). The T-DNA was inserted 2674-bp upstream of the start codon of Mapoly0060s0052.1 (Mp*ERF13*), which encodes a putative ERF/AP2 transcription factor in the subgroup containing DREB1A and TINY in Arabidopsis (Fig. S4B). Moreover, Mp*ERF13* and Mp*SYP12B* transcripts accumulated in this mutant compared to wild type as detected by RNA-sequencing (RNA-Seq) and quantitative RT-PCR analyses (Fig. S5B). We then generated knockout mutants in which the Mp*ERF13* gene was deleted by genome editing (Fig. S4C). Two independent mutants (Mp*erf13-1*^*ge*^ and Mp*erf13-2*^*ge*^) exhibited no detectable abnormalities in thallus development and reproductive growth; however, these mutants completely lacked oil bodies in the gemma and thallus tissues (Figs. 5C-E and S4F, G). These phenotypes of gain-and loss-of-function mutations suggested that MpERF13 is a major transcription factor regulating oil body formation. Consistently, the Mp*ERF13* promoter was highly active in oil body cells (Fig. 5F).

**Fig. 5.**
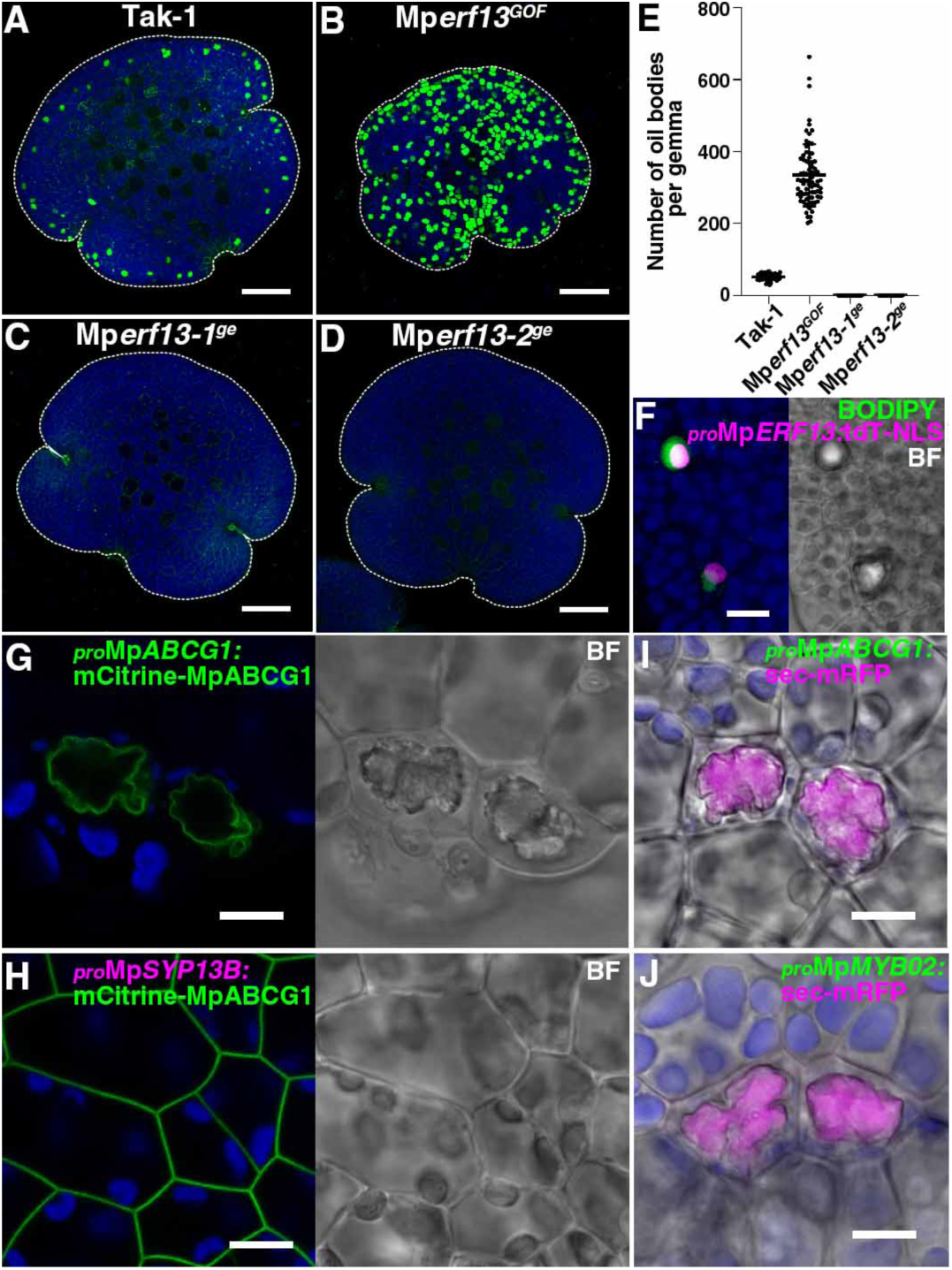
MpERF13 transcription factor regulates oil body formation. (A - D) Maximum - intensity projection images of BODIPY-stained gemmae of Tak-1 (A), Mp*erf13*^*GOF*^ (B), Mp*erf13-1*^*ge*^ (C), and Mp*erf13-2*^*ge*^ (D). (E) The number of oil bodies visualized by BODIPY-staining in gemmae. Bars indicate means ± SD. Statistical analyses between Tak-1 and each genotype were conducted using a two-tailed Welch’s *t*-test. Sample numbers were 57 gemmae for Tak-1, 78 for Mp*erf13*^*GOF*^, 61 for Mp*erf13-1*^*ge*^, and 75 for Mp*erf13-2*^*ge*^; and *p*-values are 1.19×10^−43^ for Mp*erf13*^*GOF*^, 1.44×10^−45^ for Mp*erf13-1*^*ge*^, and 1.44×10^−45^ for Mp*erf13-2*^*ge*^. (F) A maximum intensity projection image of oil body and non-oil body cells in a gemma expressing tandemTomato (tdT)-NLS driven by the Mp*ERF13* promoter. (G and H) Thallus cells including oil body cells expressing mCitrine-MpABCG1 under its own regulatory elements (G) and the Mp*SYP13B* promoter (H). BF, bright field images. (I and J) Thallus cells expressing sec-mRFP under the Mp*ABCG1* (I) and the Mp*MYB02* (J) promoters. Bright field images are merged. Green, magenta, and blue pseudo-colors indicate fluorescence from BODIPY or mCitrine, tdT or mRFP, and chlorophyll, respectively. Bars = 100 µm in (A - D) and 10 µm in (F - J).

As MpERF13 is homologous to ERF/AP2 transcription factors, this protein was expected to be involved in the transcriptional regulation of oil body formation and/or the oil body cycle. To identify genes downstream of MpERF13, we analyzed transcriptomes in Tak-1 (wild type) and Mp*erf13*^*GOF*^ and Mp*erf13-1*^*ge*^ mutant lines by RNA-Seq. Through the comparison between the three groups using an ANOVA-like test, we identified 136 differentially-expressed genes (DEGs) other than Mp*ERF13* whose expression was higher than two log_2_-fold change (FC) in Mp*erf13*^*GOF*^ compared with Tak-1, and lower than −2 log_2_FC in Mp*erf13-1*^*ge*^ than Tak-1 (FDR < 0.01) (Supplementary Table S1). To verify the RNA-Seq result, we also performed the quantitative RT-PCR analysis for selected 11 DEGs including Mp*ERF13* and Mp*SYP12B*, which confirmed that all of these DEGs were highly expressed in Mp*erf13*^*GOF*^, but their expression was significantly lower or not detectable in Mp*erf13-1*^*ge*^ and Mp*erf13-2*^*ge*^ compared with Tak-1 (Fig S5). These data strongly suggested that these 136 genes function downstream of Mp*ERF13* in oil body cells, which was further verified for the two genes Mapoly0083s0014.1 and MpMYB02 (Mapoly0006s0226). Mapoly0083s0014.1 was positively regulated by MpERF13 and encodes a member of the G subfamily of ABC transporters ^20^, which we named MpABCG1. Although ABCG members are generally targeted to the PM and mediate efflux of secondary metabolites from intracellular to extracellular spaces ^27,28^, mCitrine-MpABCG1 was targeted to the oil body membrane when expressed under its own regulatory elements (Fig. 5G); however, this protein was targeted to the PM in non-oil body cells when expressed under the Mp*SYP13B* promoter (Fig. 5H). MpMYB02, whose expression was also positively regulated by MpERF13, is reported to be a transcription factor responsible for the production of marchantin A and its derivatives that accumulate in oil bodies ^11,29^. Mp*ABCG1* and Mp*MYB02* are specifically expressed in oil body cells and when sec-mRFP was driven by the promoters of these genes, it was specifically targeted to the lumen of oil bodies. These results indicated that MpABCG1 and MpMYB02, presumably together with other DEGs we identified, act during oil body formation downstream of MpERF13.

Lastly, we asked why liverworts possess oil bodies. Using fresh and alcohol-washed liverworts, the oil body was proposed to be an effective chemical protection from herbivores ^30^, although its true function remains unknown because alcohol can wash out substances as well as oil body constituents. We fed starved *Armadillidium vulgare* (pill bug) with wild-type liverwort and genetically established mutants that contain extra or no oil bodies. Thalli of Tak-1 and Mp*erf13*^*GOF*^ remained almost intact after 24 h of herbivory, while the areas of Mp*erf13-1*^*ge*^ and Mp*erf13-2*^*ge*^ thalli were significantly reduced after herbivory (Figs. 6 and S6). This result demonstrated that the oil body is effective in protecting liverworts from herbivores.

**Fig. 6.**
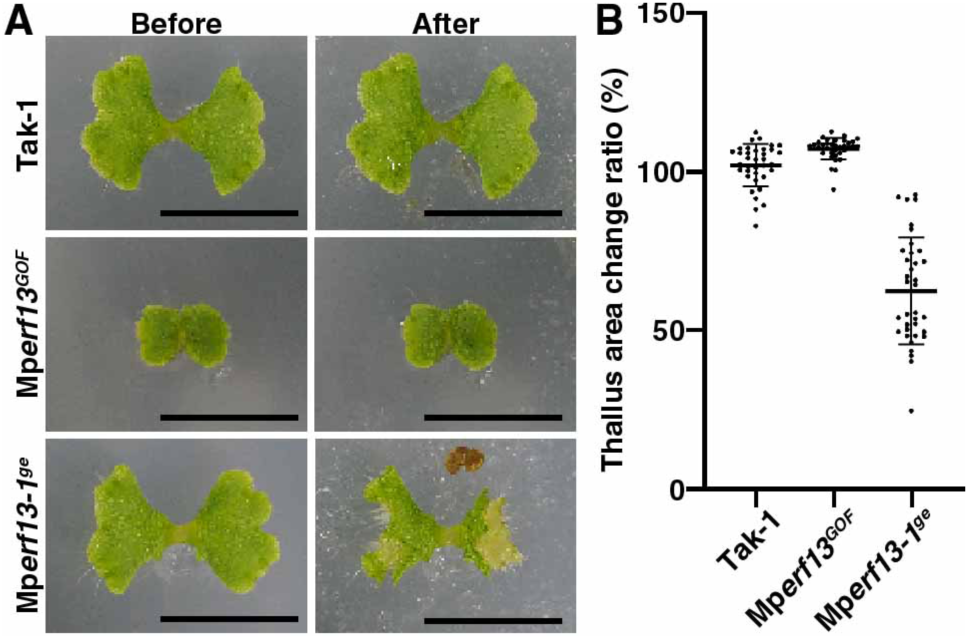
(A) Ten-day-old thalli in indicated genotypes (Before) they were fed to starved pill bugs or 24 h after feeding (After). (B) Quantification of fluorescence signals images in panel (A). Ratios of thallus areas of “After” to “Before” pill bug feeding were calculated to indicate thallus area change ratios (n = 36 thalli for each genotype). Bars indicate means ± SD. Statistical analyses between Tak-1 and each genotype were conducted by two-tailed Welch’s *t*-test. The *p*-values were 5.21×10^−4^ for Mp*erf13*^*GOF*^ and 1.05×10^−16^ for Mp*erf13-1*^*ge*^. Bars = 1 cm.

We demonstrate that plants have acquired at least two organelles, the cell plate and oil body, through a common evolutionary pathway of redirecting secretory patterns from an outward to an inward direction in cells. The functions of these organelles are totally different but they share several important traits that include specific SYP11/12 members that reside at their membranes, clathrin-coated vesicles generated from the plasma and internal membranes, and both the cell plate and oil body are formed by reorientation of secretory pathways. In both cases, orientation of the secretory pathway is switched in response to transitioning cellular states, during the cell cycle for cell plate formation and the oil body cycle for oil body formation. Orientation of the plant secretory pathway is also modulated during plant-microbe interactions; symbiotic arbuscular mycorrhizal fungi redirects the plant secretory pathway to form the arbuscule in host plant cells that acts as an interface for sugar and mineral exchange ^31,32^. Thus, redirection of the secretory pathway is a common strategy for plants to acquire novel organelles/cellular structures that had resulted in maximizing fitness for land plants during evolution. The liverwort oil body is a nice system to evaluate how novel organelles have been acquired during evolution, and the knowledge obtained from this study is useful for engineering new organelles/cellular structures in plants to maximize plant cellular functions and provides a powerful methodology for future organelle technology.

## Acknowledgments

We thank Dr. Ikuko Hara-Nishimura (Konan University), Dr. Kimitsune Ishizaki (Kobe University), Dr. Ryuichi Nishihama (Kyoto University), and Dr. Shigeo S. Sugano for sharing vectors and plant materials, Ms. Miwako Matsumoto, Ms. Asaka Akita, and Functional Genomics Facility (NIBB) for technical assistance in RNA-Seq experiments, and Dr. Ken Naito (National Agriculture and Food Research Organization) for instruction for RNA-Seq data analysis. We thank Dr. John L. Bowman (Monash University) and Dr. Facundo Romani (Universidad Nacional del Litoral) for helpful discussion. We also thank Dr. Teruyuki Niimi (NIBB) and Dr. Takahisa Miyatake (Okayama University) for identification of the species of pill bug used in this study. The support of plant cultivation rooms was provided by the Model Plant Research Facility of NIBB.

## Author contributions

T.Ka. performed the majority of the experiments, H.M. performed the experiment presented in Fig. 1A-G, K.E. and T.L.S. provided essential ideas for experiments using sec-mRFP targeting and oil body detection, respectively, S.I. generated the Mp*syp12b-1* mutant, K.Y. and S.S. designed the RNA-seq experiment, N.M. examined the effect of Mp*erf13* mutations on reproductive growth, T.Ka. and T.U. wrote the manuscript, and T.Ko., A.N., and T.U. supervised the study.

## Data and materials availability

All reads of RNA-Seq are available through the Sequence Read Archive (SRA) under the accession number DRA009193.

## Supplementary Information for

### Materials and Methods

#### Phylogenetic Analyses

Phylogenetic analyses were performed as previously described in Bowman *et al*, 2017 ^20^ with minor modification. Previously published datasets in Kanazawa *et al*., 2016 and Bowman *et al*., 2017, were used for phylogenetic analyses of SYP1 and ERF/AP2 members, respectively ^13,20^. Multiple sequence alignments were performed using the MUSCLE program version 3.8.31 ^33,34^ with the default parameter. The alignment gaps were removed by Gblocks program version 0.91b ^35,36^ or manually. The maximum likelihood phylogenetic analyses were performed using PhyML 3.0 ^37^ under the LG model. Bootstrap analyses were performed by resampling 1000 sets.

#### Vector Construction

Open reading frames (ORFs) and genomic sequences of Marchantia genes were amplified by polymerase chain reaction (PCR) from cDNA and genomic DNA prepared from the accession Tak-1, and the amplified products were subcloned into pENTR/D-TOPO (ThermoFisher, Waltham, USA) according to the manufacturer’s instructions. The primer sequences and sizes of amplified products are listed in Supplementary Table S2. For construction of pENTR _*pro*_Mp*ERF13*, the 5′ sequence (promoter + 5′ UTR) was amplified and subcloned into pENTR/D-TOPO. The resultant sequence was introduced into pMpGWB316 ^38^ using the Gateway LR Clonase™ II Enzyme Mix (ThermoFisher) according to the manufacturer’s instructions. For construction of pENTR genomic mGFP-MpSYP12A and mRFP-MpSYP13B constructs, the cDNA for mGFP and mRFP were inserted into *Sma*I and *Bam*HI sites in the pENTR genomic XFP(*Sma*I)-MpSYP12A and pENTR genomic XFP(*Bam*HI)-MpSYP13B, which were previously prepared ^13^ using the In-Fusion HD Cloning System (Clontech, Shiga, Japan). For construction of pENTR genomic mCitrine-MpPIP2, pENTR genomic mCitrine-MpSYP4, pENTR genomic mCitrine-MpSEC22, and pENTR genomic mCitrine-MpABCG1, genomic sequences of the protein-coding regions and 3′ flanking sequences were amplified with a *Sma*I site at the 5′ end and subcloned into the pENTR vector. The 5′ sequences (promoter + 5′ UTR) and cDNA of mCitrine were amplified and inserted into *Not*I and *Sma*I sites, respectively, of the resultant pENTR vectors using the In-Fusion HD Cloning System. The resultant chimeric genes were then introduced into pMpGWB301 ^38^ using the Gateway LR Clonase™ II Enzyme Mix. For construction of pENTR _*pro*_Mp*SYP12B:mCitrine*, the _*pro*_Mp*SYP12B:mCitrine* sequence was amplified from the pENTR genomic mCitrine-MpSYP12B vector, and subcloned into pENTR/D-TOPO. The resultant sequence was introduced into pMpGWB307 ^38^ using the Gateway LR Clonase™ II Enzyme Mix to create the pMpGWB307 _*pro*_Mp*SYP12B:2*×*YFP* construct. For construction of pENTR genomic MpCLC1-mCitrine, genomic sequence comprising the 5′ flanking (promoter + 5′ UTR) and protein-coding sequences were amplified with a *Sma*I site at the 3′ end and subcloned into the pENTR vector. The 3′ flanking sequence including 3′ UTR and cDNA for mCitrine were amplified and inserted into *Asc*I and *Sma*I sites, respectively, of the resultant pENTR vectors using the In-Fusion HD Cloning System. The resultant chimeric genes were then introduced into pMpGWB301 ^38^ using the Gateway LR Clonase™ II Enzyme Mix. For construction of pENTR sec-mRFP, sequences for the signal peptide and mRFP were independently amplified by PCR, and the sequence of sec-mRFP was then amplified by PCR using the mixture of the amplified products as templates. The amplified products were subcloned into pENTR/D-TOPO. For the construction of pENTR mRFP-HDEL, the sequence of SP-mRFP-HDEL was amplified by PCR using pENTR sec-mRFP as template, and subcloned into pENTR/D-TOPO. For construction of pENTR _*pro*_Mp*SYP12B:*sec-mRFP, pENTR _*pro*_Mp*SYP13B:*sec-mRFP, pENTR _*pro*_Mp*ABCG1:*sec-mRFP, and pENTR _*pro*_Mp*MYB02:*sec-mRFP, the 5′ sequences (promoter + 5′ UTR) were amplified and were inserted into *Not*I sites of the pENTR sec-mRFP vector using the In-Fusion HD Cloning System. The resultant sequences were then introduced into pMpGWB301 ^38^ using the Gateway LR Clonase™ II Enzyme Mix. To construct pMpGWB301-derived Gateway vectors (Fig. S2A), the promoter:*mCitrine* sequences were amplified from genomic constructs described above, and inserted at a *Hin*dIII site of pMpGWB301 using the In-Fusion HD Cloning System. pENTR MpTUB2 was kindly provided from Dr. R. Nishihama and introduced into pMpGWB301_*pro*_Mp*SYP2:mCitrine*-GW (Fig. S2A) using the Gateway LR Clonase™ II Enzyme Mix. For construction of the genome-editing vectors, the target sequences were selected using CRISPR direct (https://crispr.dbcls.jp/) ^39^, and double-stranded oligonucleotides of the target sequences were inserted into the pMpGE_En03 vector ^40^. The resultant gRNA cassettes were introduced into the pMpGE010 or pMpGE011 vectors ^40^ using the Gateway LR Clonase II Enzyme Mix. For construction of the homologous recombination-mediated gene targeting vector, the 3.5-kb homologous genomic sequences were amplified from the Tak-1 genome and inserted at *Pac*I and *Asc*I sites of the pJHY-TMp1 vector ^41^ using the In-Fusion HD Cloning System.

#### Plant materials and Transformation

Marchantia accession Takaragaike-1 (Tak-1, male) and Takaragaike-2 (Tak-2, female) ^42^ were used throughout the study. The growth conditions and transformation methods were previously described ^13,42,43^. *M. polymorpha* expressing mCitrine-MpSYP12A, mCitrine-MpSYP12B, mCitrine-MpSYP13A, mCitrine-MpSYP13B, mCitrine-MpSYP2 or mCitrine-MpVAMP71 under the regulation of their own regulatory elements (e.g. promoter, UTRs, and introns), and *M. polymorpha* expressing mRFP-MpSYP6A or GFP-HDEL driven by the Mp*EF1α* promoter were previously reported ^13,38,44^.

#### Genotyping

Genomic DNA was extracted from the thalli in an extraction buffer containing 1 M KCl, 100 mM Tris-HCl (pH 9.5), and 10 mM EDTA. This DNA was used as templates for PCR. Genome fragments with putative mutation sites were amplified by PCR using KOD FX Neo (TOYOBO, Osaka, Japan) according to the manufacturer’s instructions. The primers used in PCR-based genotyping are listed in Supplementary Table S2.

#### Confocal laser scanning microscopy

Two- and five-day-old thalli were used for imaging the cell plate and the oil body, respectively. Samples were mounted in distilled water and observed using an LSM780 confocal microscope (Carl Zeiss) equipped with an oil immersion lens (63×, numerical aperture = 1.4) on an electrically driven stage. Gemmae were observed with a 10× objective lens (numerical aperture = 0.45). Thalli were incubated in 10 µM *N*-(3-triethylammoniumpropyl)-4-(6-(4-(diethylamino)phenyl)hexatrienyl) pyridinium dibromide (FM4-64, Thermo Fisher) solution for 2 min. Samples were then washed twice in water before observation. For 4,4-difluoro-1,3,5,7,8-pentamethyl-4-bora-3a,4a-diaza-*s*-indacene (BODIPY 493/503, Thermo Fisher) staining, thalli or gemmae were incubated in 200 nM BODIPY 493/503 dissolved in water for 10 min and then washed twice in water before observation. The samples were excited at 488 nm (Argon 488) and 561 nm (DPSS 561-10) lasers, respectively, and the fluorescent emission was collected between 482-659 nm using twenty GaAsP detectors. Spectral unmixing and constructing maximum intensity projection images were conducted using the ZEN2012 software (Carl Zeiss). Image processing was performed with ZEN2012 software, Photoshop software (Adobe Systems), and ImageJ (National Institute of Health, https://imagej.nih.gov/ij/).

#### Electron microscopy

Five-day-old *M. polymorpha* thalli from Tak-1 and thalli expressing mCitrine-MpSYP12B were used for the immunoelectron microscopic observation. Sample preparation and observation were performed as previously reported ^13^. Five-day-old Tak-1 thalli were used for morphological observation with a transmission electron microscope. Sample preparation and observation were performed as previously reported ^44^.

#### Light sheet microscopy

Five-day-old thalli expressing 2×Citrine driven by the Mp*SYP12B* promoter were embedded in low melt agarose gel and observed using a Light sheet Z.1 microscope (Carl Zeiss) equipped with a water immersion lens (5×, numerical aperture = 0.16). The samples were excited at 488 nm (Argon 488 laser). Acquisition and construction of three-dimension images from multi-angle images were conducted using the ZEN2014 software (Carl Zeiss). The images were processed digitally with Imaris (Bitplane) and Photoshop software.

#### Fluorescent stereoscopic microscopy

Three-week-old wild-type and Mp*erf13*^*ge*^ thalli were stained by BODIPY493/503 (ThermoFisher) and observed using a fluorescent stereoscopic microscope (M165 FC, Leica) equipped with a MC170 HD digital camera (Leica).

#### Forward genetic screening for the oil body formation mutants

The workflow of the mutant screening is described in Fig. S4A. Liverworts with transfer DNA (T-DNA) insertions were generated by co-culture of *M. polymorpha* sporelings with the agrobacteria strain GV2260 harbouring the binary vector pCAMBIA1300 (https://cambia.org/welcome-to-cambialabs/cambialabs-projects/cambialabs-projects-legacy-pcambia-vectors-pcambia-legacy-vectors-1/cambialabs-projects-legacy-pcambia-vectors-list-of-legacy-pcambia-vectors-3/). T_1_ plants were first selected on 1/2× Gamborg’s B5 plates containing 10 mgL^-1^ hygromycin and 100 mgL^-1^ cefotaxime. Hygromycin-resistant T_1_ plants were then transferred and incubated on hygromycin-free medium for about one week, followed by BODIPY 493/503 staining to visualize oil bodies. Visible screening for mutants defective in oil body formation was performed using a fluorescent stereoscopic microscope (M165 FC, Leica). To identify flanking sequences of T-DNA insertions, thermal asymmetric interlaced-PCR (TAIL-PCR) was performed as described previously ^42,45,46^ with minor modifications using crude-extracted DNA as templates. Flanking sequences were amplified using KOD FX neo DNA polymerase and T-DNA-specific (TR1–3 and TL1–3 for right and left borders of T-DNA, respectively) and universal adaptor (AD1–6) primers. The reaction cycles are shown below. After agarose gel electrophoresis of the final TAIL-PCR products, DNA bands were excised and purified using the Wizard SV Gel and PCR Clean-Up System (Promega). Purified products were directly sequenced using TR3 or TL3 primers. The T-DNA insertion sites were identified using the genome sequence registered in MarpolBase, genome version 3.1.

#### Cycling conditions for TAIL-PCR

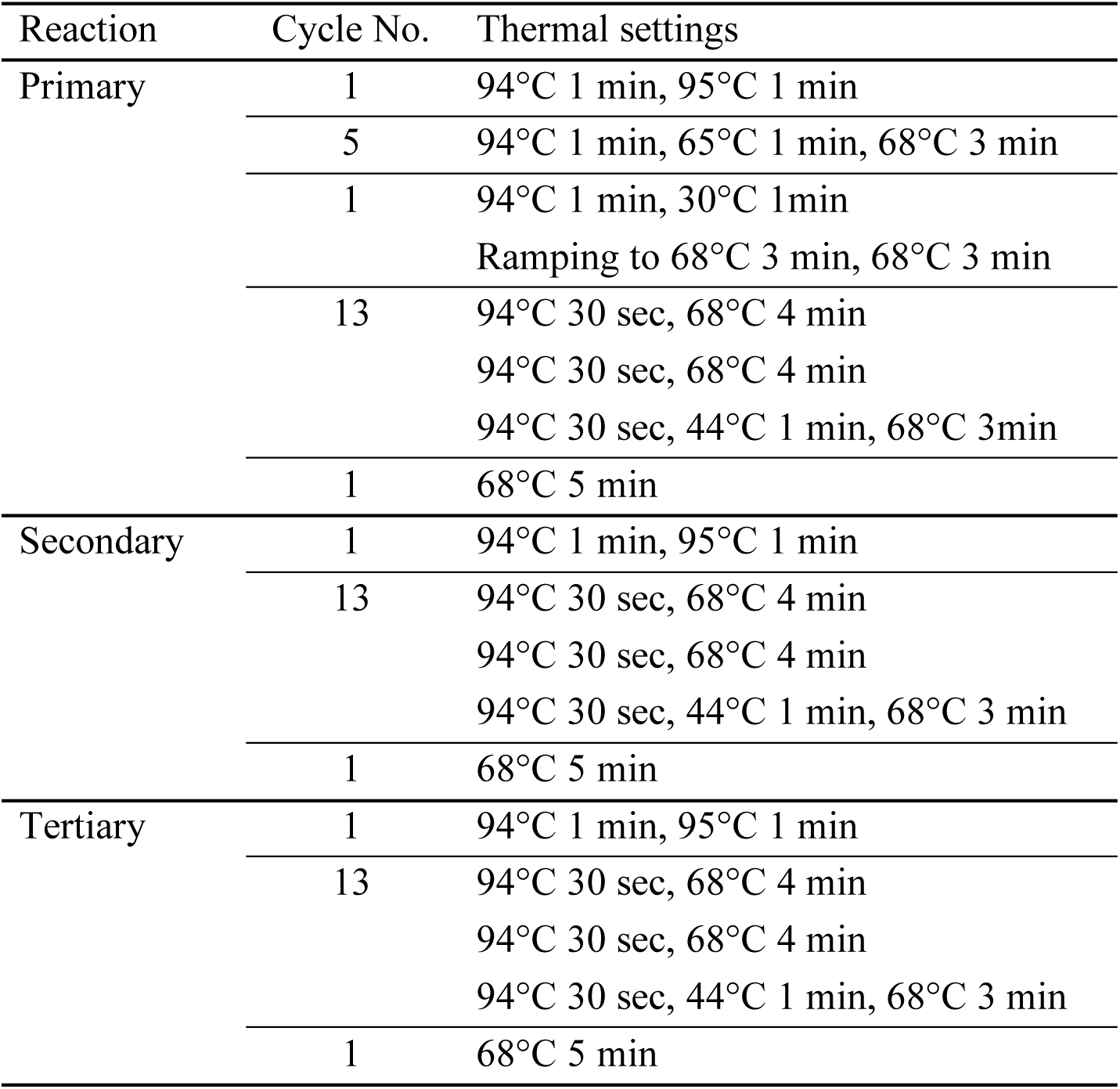

#### RNA extraction and RNA sequencing analysis

Total RNA from Tak-1, Mp*erf13*^*GOF*^ and Mp*erf13-1*^*ge*^ thalli was extracted by RNeasy (QIAGEN) according to the manufacturer’s instructions. The eluted total RNA samples were treated with DNase I (Takara) to remove DNA contamination. The quality and quantity of the total RNA were evaluated with a NanoDrop 1000 spectrophotometer (Thermo Fisher Scientific), Qubit 2.0 Fluorometer (Thermo Fisher Scientific) and a Bioanalyzer RNA6000 Nano Kit (Agilent Technologies). The sequence libraries were prepared with the TruSeq RNA Sample Prep Kit v2 (Illumina) according to the manufacturer’s low sample protocol. The quality and quantity of each library were determined using a Bioanalyzer with the High Sensitivity DNA kit (Agilent Technologies) and the KAPA Library Quantification Kit for Illumina (Illumina). Equal amounts of each library were mixed to make the 2 nM pooled library. Illumina sequencing was performed using a HiSeq 1500 platform (Illumina) to produce 126-bp single-end reads. Three biological replicates were prepared for the library construction and RNA-Seq analysis. All reads are available through the Sequence Read Archive (SRA) under the accession number DRA009193. The abundance of transcripts from Illumina RNA-Seq data were quantified with the kallisto program (version 0.43.1) ^47^ with the default parameters using the primary transcript dataset from the Marchantia genome version 3.1. To estimate differentially expressed genes (DEGs), an ANOVA-like test was performed by edgeR (version 3.18.1) ^48^. Genes with FDR values lower than 0.01 and an absolute log_2_ fold change (FC) > 2 were considered to be differentially expressed. Among all combinations of DEGs (log_2_FC(Mp*erf13*^*GOF*^/Tak-1) > 2 and log_2_FC(Mp*erf13-1*^*ge*^/Tak-1) > 2, log_2_FC(Mp*erf13*^*GOF*^/Tak-1) > 2 and log_2_FC(Mp*erf13-1*^*ge*^/Tak-1) < −2, log_2_FC(Mp*erf13*^*GOF*^/Tak-1) < −2 and log_2_FC(Mp*erf13-1*^*ge*^/Tak-1) > 2, log_2_FC(Mp*erf13*^*GOF*^/Tak-1) < −2 and log_2_FC(Mp*erf13-1*^*ge*^/Tak-1) < −2), we selected the set log_2_FC(Mp*erf13*^*GOF*^/Tak-1) > 2 and log_2_FC(Mp*erf13-1*^*ge*^/Tak-1) < −2 as candidate genes regulated downstream of MpERF13, which included both Mp*ERF13* and Mp*SYP12B*.

#### Quantitative reverse transcription (qRT)-PCR

For qRT-PCR, cDNA was synthesized from total RNA using SuperScript III Reverse Transcriptase (Invitrogen) and an oligo dT(18) primer according to the manufacturer’s instructions. qRT-PCR was performed with LightCycler 480 (Roche) using FastStart SYBR Green Master (Roche) according to the manufacturer’s protocol. Sequences of primers used are listed in Table S2. Mp*APT* (Mapoly0100s0027.1 / Mp3g25140.1) was used as a housekeeping reference for normalization ^49^. Three biological replicates were prepared and three technical replicates were performed for each reaction.

#### Pill bug feeding assay

Feeding assay was performed according to Nakazaki *et al*. with modification ^50^. Gemmae were cultured on 1/2× Gamborg’s B5 medium containing 1% (w/v) agar and 1% (w/v) sucrose for five days at 22°C under continuous white light (50 µmol m^-2^s^-1^). Five-day-old thalli were transferred onto 1/2× Gamborg’s B5 medium containing 1% (w/v) agar without sucrose and cultured for additional five days under the same conditions. Pill bugs (*Armadillidium vulgare*) were collected in the Myodaiji area of NIBB (Nishigonaka 38, Myodaiji, Okazaki 444-8585 Aichi, Japan). Pill bugs were maintained on Prowipe (Daio Paper, Tokyo, Japan) moistened with sterilized water for 48 h at 22°C in the dark without feeding before the assay. Six pill bugs were introduced into each medium plate containing 10-day-old thalli and kept for 24 h in the dark at 22°C. The thallus areas were calculated using ImageJ (National Institute of Health, https://imagej.nih.gov/ij/).

#### Gene accession numbers

| Symbol | Gene ID (ver3.1) | Gene ID (ver5.1) |
| --- | --- | --- |
| Mp*SYP12A* | Mapoly0163s0015 | Mp6g00050 |
| Mp*SYP12B* | Mapoly0101s0013 | Mp4g20670 |
| Mp*SYP13A* | Mapoly0009s0005 | Mp7g13190 |
| Mp*SYP13B* | Mapoly0055s0091 | Mp2g19600 |
| Mp*CLC1* | Mapoly0075s0069 | Mp2g03080 |
| Mp*PIP2* | Mapoly0041s0003 | Mp4g17210 |
| Mp*ERF13* | Mapoly0060s0052 | Mp6g08690 |
| Mp*ABCG1* | Mapoly0083s0014 | Mp8g13070 |
| Mp*MYB02* | Mapoly0006s0226 | Mp3g07510 |
| Mp*TUB2* | Mapoly0158s0010 <sup>51,52</sup> | Mp2g09390 |
| Mp*SYP6A* | Mapoly0140s0004 | Mp3g18380 |
| Mp*SYP2* | Mapoly0187s0013 | Mp8g15260 |
| Mp*SFT1* | Mapoly0006s0109 | Mp3g06390 |
| Mp*GOS11* | Mapoly0001s0245 | Mp1g19070 |
| Mp*SYP4* | Mapoly0042s0041 | Mp2g14140 |
| Mp*VAMP71* | Mapoly0064s0109 | Mp8g00880 |
| Mp*SEC20* | Mapoly0019s0137 | Mp1g13670 |
| Mp*USE1A* | Mapoly0001s0213 | Mp1g18750 |
| Mp*SEC22* | Mapoly0023s0071 | Mp2g11050 |
| Mp*CYCB1* | Mapoly0048s0068 | Mp5g10030 |

We followed the nomenclature of genes, proteins, and mutants of Marchantia reported in Bowman *et al*. (2016) ^53^. Gene IDs were taken from MarpolBase (http://marchantia.info/), genome version 3.1 and version 5.1 ^20,54^.

**Fig. S1.**
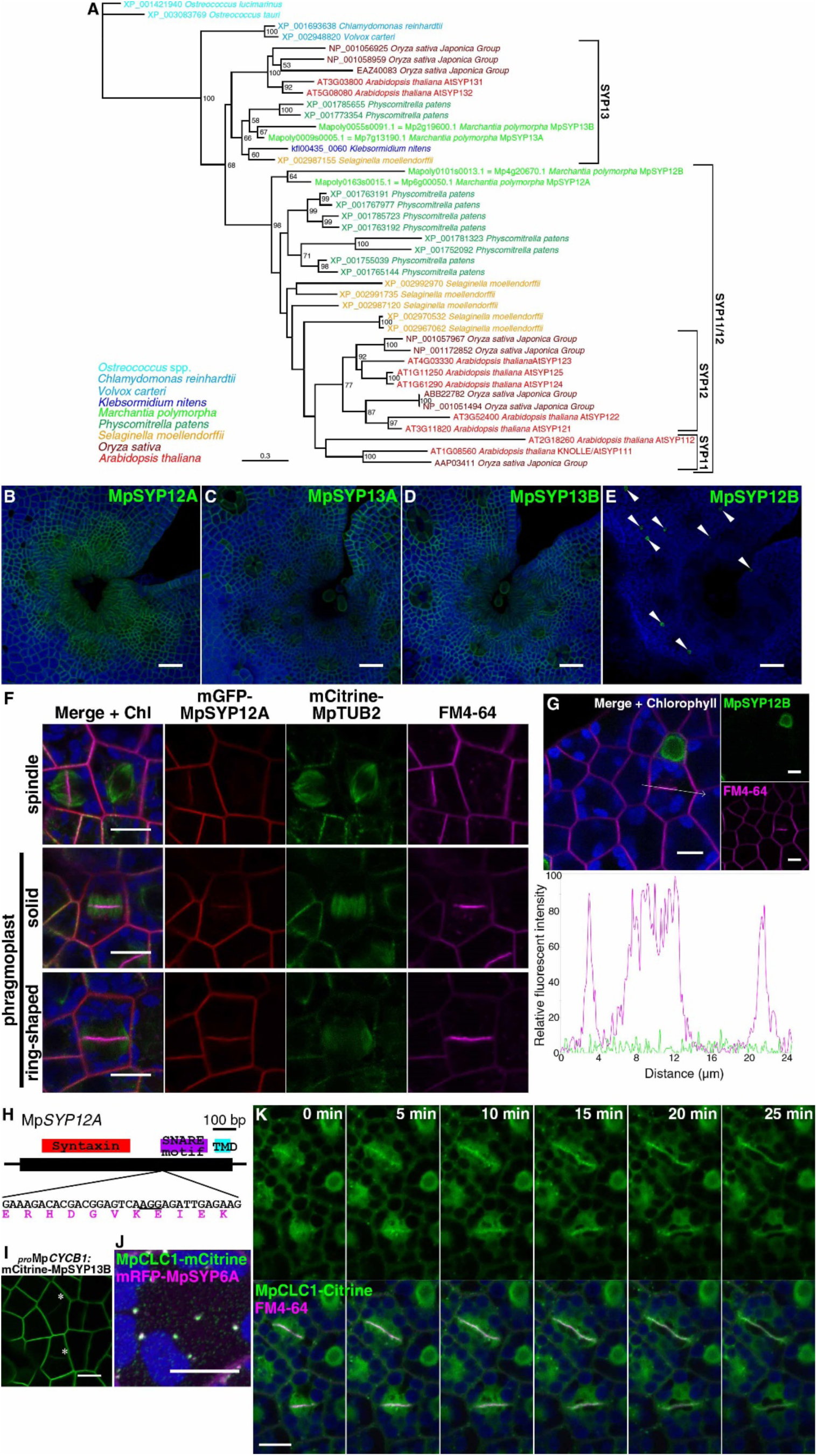
Characterization of SYP1 member and clathrin light chain localization during cytokinesis in Marchantia thalli. (A) A maximum likelihood phylogenetic tree of SYP1 members in green plants. Although the tree is unrooted, the proteins from Chlorophytes (*Ostreococcus* spp., *Chlamydomonas reinhardtii*, and *Volvox carteri*) are the outgroup to those of streptophytes. The branch lengths are proportional to the estimated number of substitutions per site. Bootstrap probability is indicated as a percentage on each branch with at least 50% support. (B - E) Maximum intensity projection images of thalli expressing mCitrine-MpSYP12A (B), mCitrine-MpSYP13A (C), mCitrine-MpSYP13B (D), or mCitrine-MpSYP12B (E). White arrowheads indicate mCitrine-MpSYP12B-expressing cells. (F) Single confocal images of thallus cells expressing mCitrine-MpTUB2 and mGFP-MpSYP12A under the regulation of the Mp*SYP2* and Mp*SYP12A* promoters, respectively, stained with FM-4-64. In mitotic cells, spindles and phragmoplasts (solid and ring-shaped phases) were labelled by mCitrine-MpTUB2, and endocytosed FM-4-64 accumulated at forming cell plates. (G) Single confocal images of thallus cells expressing mCitrine-MpSYP12B stained with FM4-64. The line graph indicates the relative fluorescence intensity along the white arrow. Relative fluorescent intensity of mCitrine to fluorescence on the oil body membrane is shown for mCitrine fluorescence. (H) The gene model and sequences of MpSYP12A. The syntaxin domain, SNARE motif, and transmembrane domain (TMD) are shown above the gene model. Black and magenta letters indicate genome and translated amino acid sequences. The protospacer adjacent motif (PAM) sequence for gRNA is underlined. (I) Expression by the Mp*CYCB1* promoter results in accumulation of MpSYP13B at the forming cell plate. A single confocal image of dividing thallus cells expressing mCitrine-MpSYP13B are shown. Asterisks indicate forming cell plates. (J) A maximum intensity projection image of a thallus cell expressing MpCLC1-mCitrine and mRFP-MpSYP6A under the regulation of their own regulatory elements and the Mp*EF1α* promoter, respectively. (K) Time-lapse single confocal images of thallus cells expressing MpCLC1-Citrine under the regulation of the Mp*EF1α* promoter (upper panels) and merged images with FM4-64 signals. Autofluorescence from chlorophyll is pseudo-colored in blue. Bars = 50 µm in (B - E) and 10 µm in (F), (G), (I), (J), and (K).

**Fig. S2.**
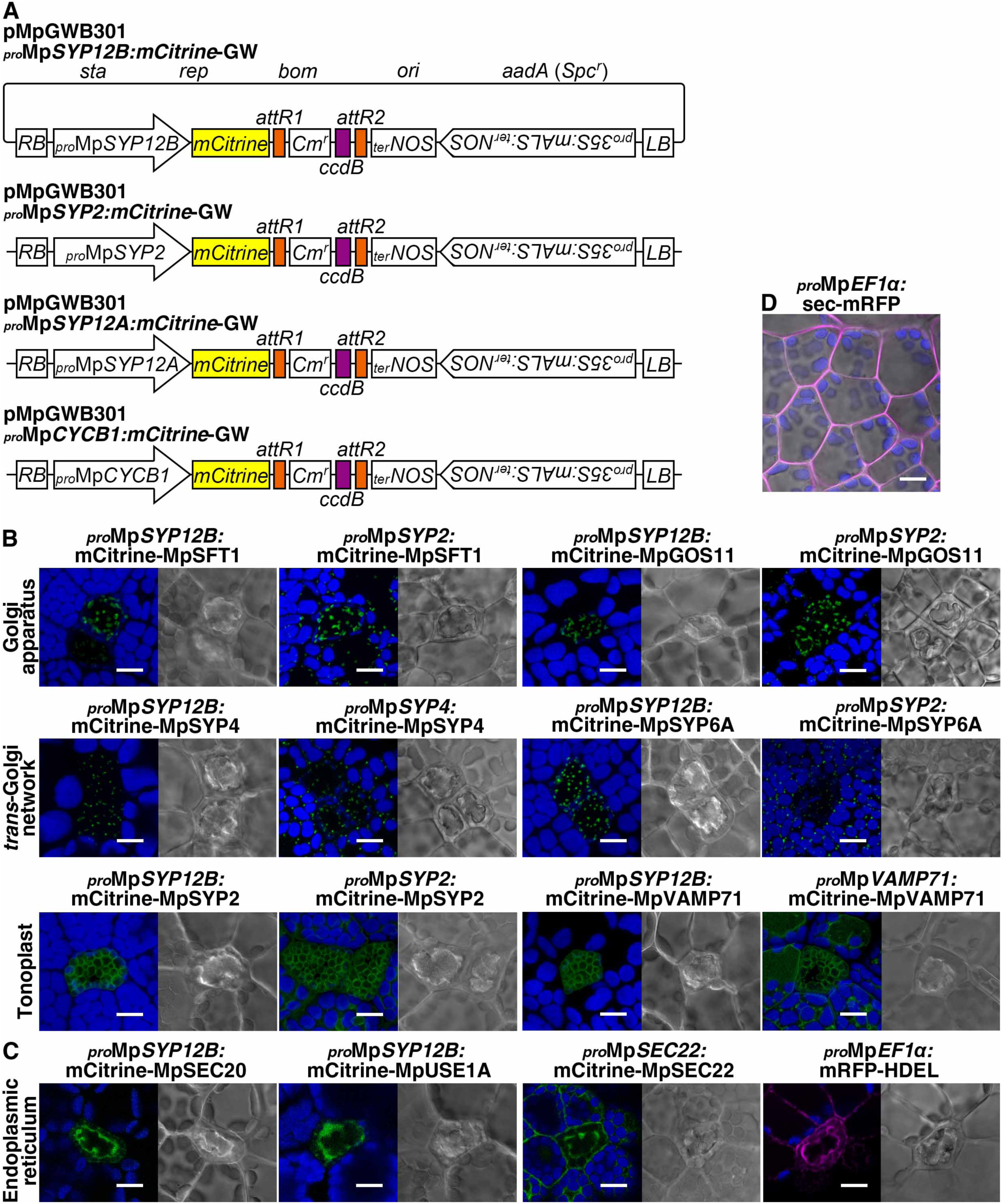
Subcellular localization of organelle markers in oil body cells. (A) Gateway constructs used in this study. pMpGWB301 _*pro*_Mp*SYP2:*mCitrine-GW was previously prepared^44^. (B) Maximum intensity projection images of thallus cells including the oil body cell expressing mCitrine-fused markers for the Golgi apparatus (MpSFT1 and MpGOS11), *trans*-Golgi network (MpSYP4 and MpSYP6A), and tonoplast (MpSYP2 and MpVAMP71). The organelle markers were expressed under the Mp*SYP12B*, Mp*SYP2*, and/or own promoters. Bright field images are also shown. (C) Single confocal images of thallus cells including the oil body cell expressing endoplasmic reticulum markers. mCitrine-MpSEC20 and mCitrine-MpUSE1A were expressed under the Mp*SYP12B* promoter. mCitrine-MpSEC22 and mRFP-HDEL were expressed using their own regulatory elements and the Mp*EF1α* promoter, respectively. (D) A single confocal image of thallus cells expressing sec-mRFP expressed under the Mp*EF1α* promoter. Green, magenta, and blue pseudo-colors indicate fluorescence from mCitrine, mRFP, and chlorophyll, respectively. Bars = 10 µm.

**Fig. S3.**
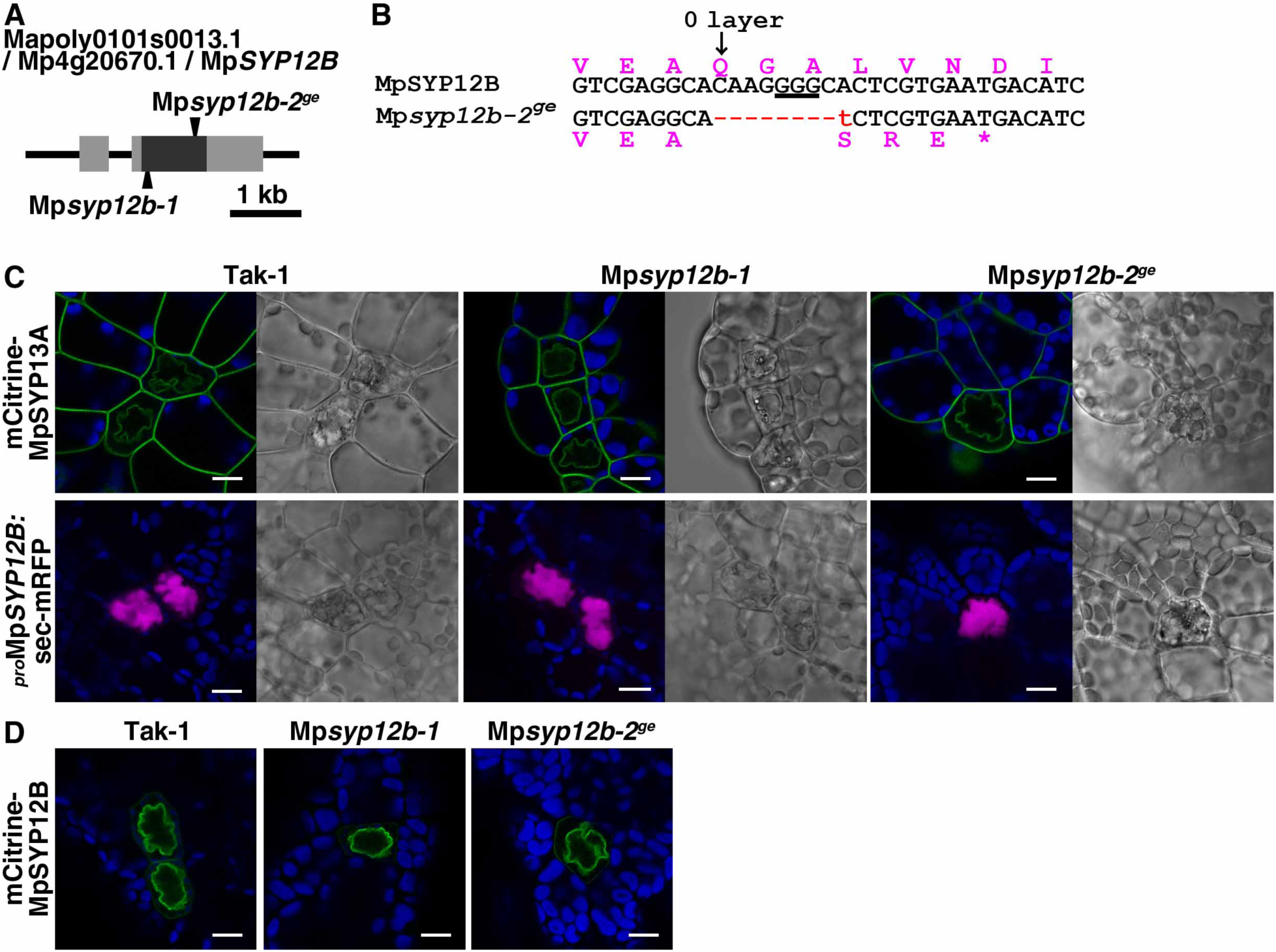
Phenotype of Mp*syp12b* mutants. (A) Schematic representation of the Mp*SYP12B* gene structure. Gray and black boxes indicate untranslated and coding regions, respectively, and arrowheads indicate mutation sites in two independently generated mutants. The Mp*syp12b-1* mutant was generated by a homologous recombination-mediated gene targeting method^41^. (B) The genome (black letters) and translated amino acid (magenta letters) sequences of Mp*SYP12B* and the Mp*syp12b-2*^*ge*^ mutant, with the red letters indicating the mutated region. The PAM sequence for gRNA is underlined. The 0 layer at the centre of the SNARE motif is shown. (C) Single confocal images of thallus cells including the oil body cell expressing mCitrine-MpSYP13A under its own regulatory elements, or sec-mRFP under the Mp*SYP12B* promoter in Tak-1, Mp*syp12b-1*, and Mp*syp12b-2*^*ge*^ lines. Bright field images are also shown. (D) Single confocal images of thallus cells including the oil body cell expressing mCitrine-MpSYP12B under its own regulatory elements in Tak-1 and Mp*syp12b* mutants. Green, magenta, and blue pseudo-colors indicate fluorescence from mCitrine, mRFP, and chlorophyll, respectively. Bars = 10 µm.

**Fig. S4.**
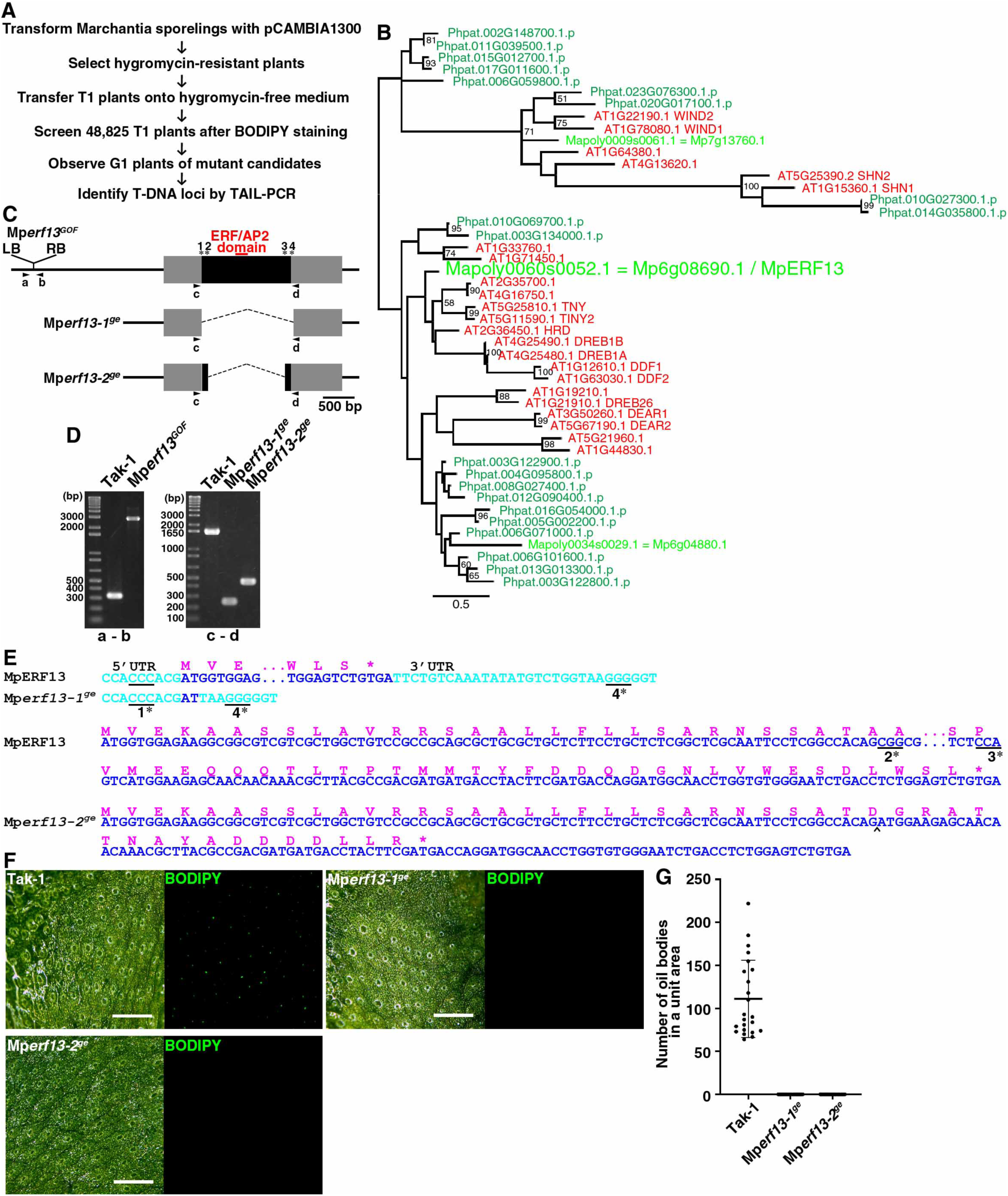
Screening and generation of Mp*erf13* mutants. (A) The flow chart of the screen for mutants defective in oil body formation from T-DNA-insertion lines. (B) A maximum likelihood phylogenetic tree of proteins with one ERF/AP2 domain. The color code is shown in Fig S1A. The branch lengths are proportional to the estimated number of substitutions per site. Bootstrap probability is indicated as a percentage on each branch with at least 50% support. A more detailed tree was presented previously (*20*). (C) Schematic representation of the Mp*ERF13* gene structure and mutations generated in this study. Gray and black boxes indicate the UTR and coding sequences, respectively. Asterisks with numbers indicate sites of designed gRNA to generate Mp*erf13-1*^*ge*^ (*1 and *4) and Mp*erf13-2*^*ge*^ (*2 and *3). (D) PCR-based genotyping of Tak-1, Mp*erf13*^*GOF*^, Mp*erf13-1*^*ge*^, and Mp*erf13-2*^*ge*^. The combinations and annealing sites of primers (a to d) are shown in (C). (E) The genomic and predicted amino acid sequences of the Mp*ERF13* locus in Tak-1 and Mp*erf13* mutants. Light blue, dark blue, and magenta letters indicate UTR, coding, and predicted amino acid sequences, respectively. The PAM sequences for gRNAs are underlined. The caret indicates the indel site. (F) Fluorescent and bright-field images of BODIPY-stained three-week-old thalli of Tak-1, Mp*erf13-1*^*ge*^, and Mp*erf13-2*^*ge*^. Bars = 0.5 mm. (G) The number of oil bodies visualized with BODIPY in a unit area (2.0 mm × 2.0 mm). Bars indicate means ± SD. Statistical analyses between Tak-1 and each genotype were conducted using a two-tailed Welch’s *t*-test. Sample numbers were 23 thalli for Tak-1, 24 for Mp*erf13-1*^*ge*^, and 26 for Mp*erf13-2*^*ge*^. *p*-values are 5.14×10^−11^ for Mp*erf13-1*^*ge*^ and 5.14×10^−11^ for Mp*erf13-2*^*ge*^.

**Fig. S5.**
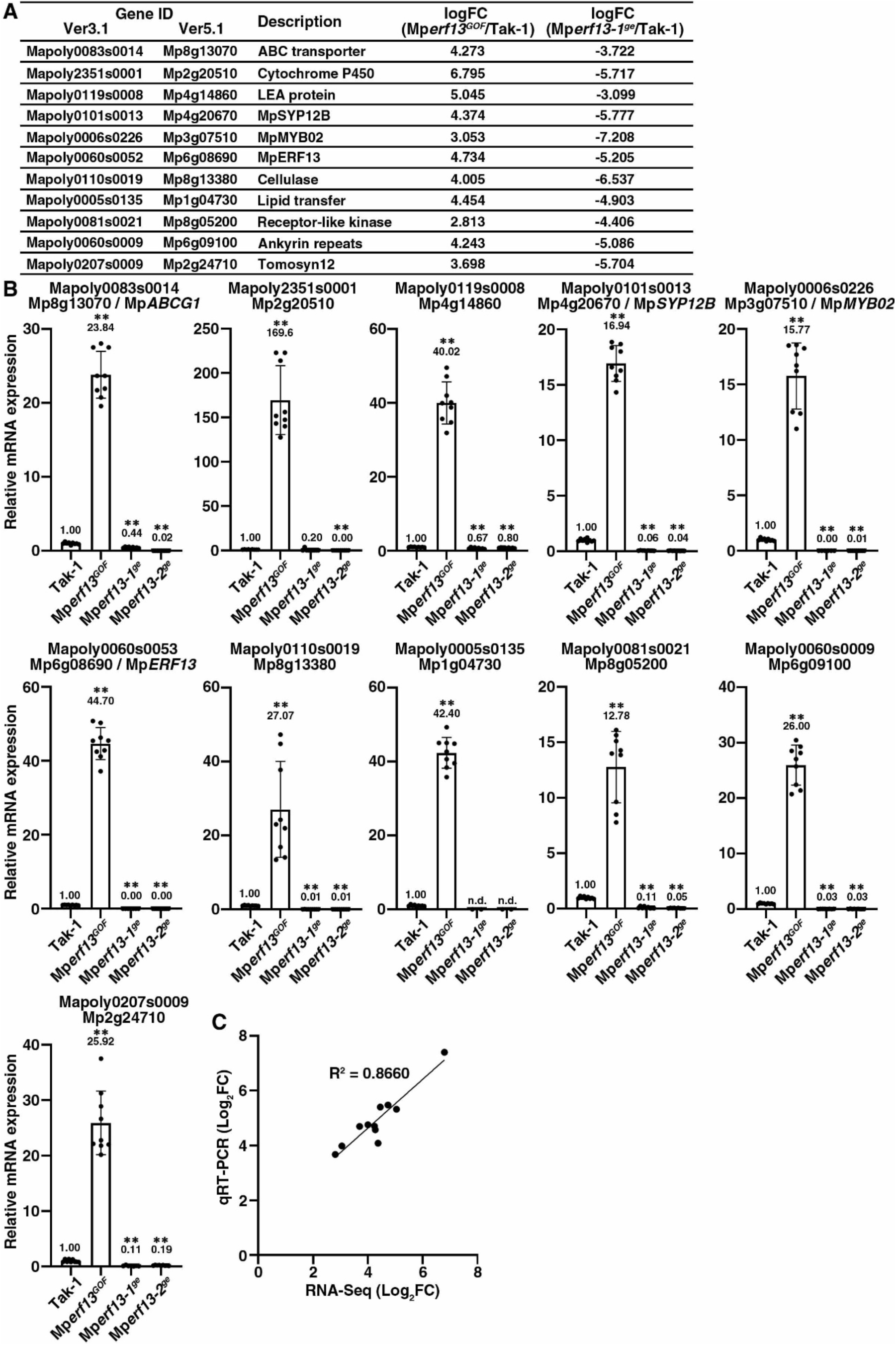
Gene expression in Mp*erf13* mutants. (A) List of selected differentially expressed genes (DEGs) for qRT-PCR analysis. The full list of DEGs are shown as Supplementary Table S1. (B) The relative mRNA expression of 11 DEGs in Tak-1, Mp*erf13*^*GOF*^, Mp*erf13-1*^*ge*^, and Mp*erf13-2*^*ge*^ measured by qRT-PCR. Mp*APT* was used as an internal reference. Error bars represent ± SD. Means are indicated above the bars. Statistical comparison between Tak-1 and each genotype was conducted with a two-tailed Welch’s *t*-test. **, *p* < 0.01. (C) Correlation of the expression levels between the RNA-Seq analysis and qRT-PCR experiment. Both the x- and y-axes are shown in a log_2_FC scale. R^2^ indicates the correlation coefficient.

**Fig. S6.**
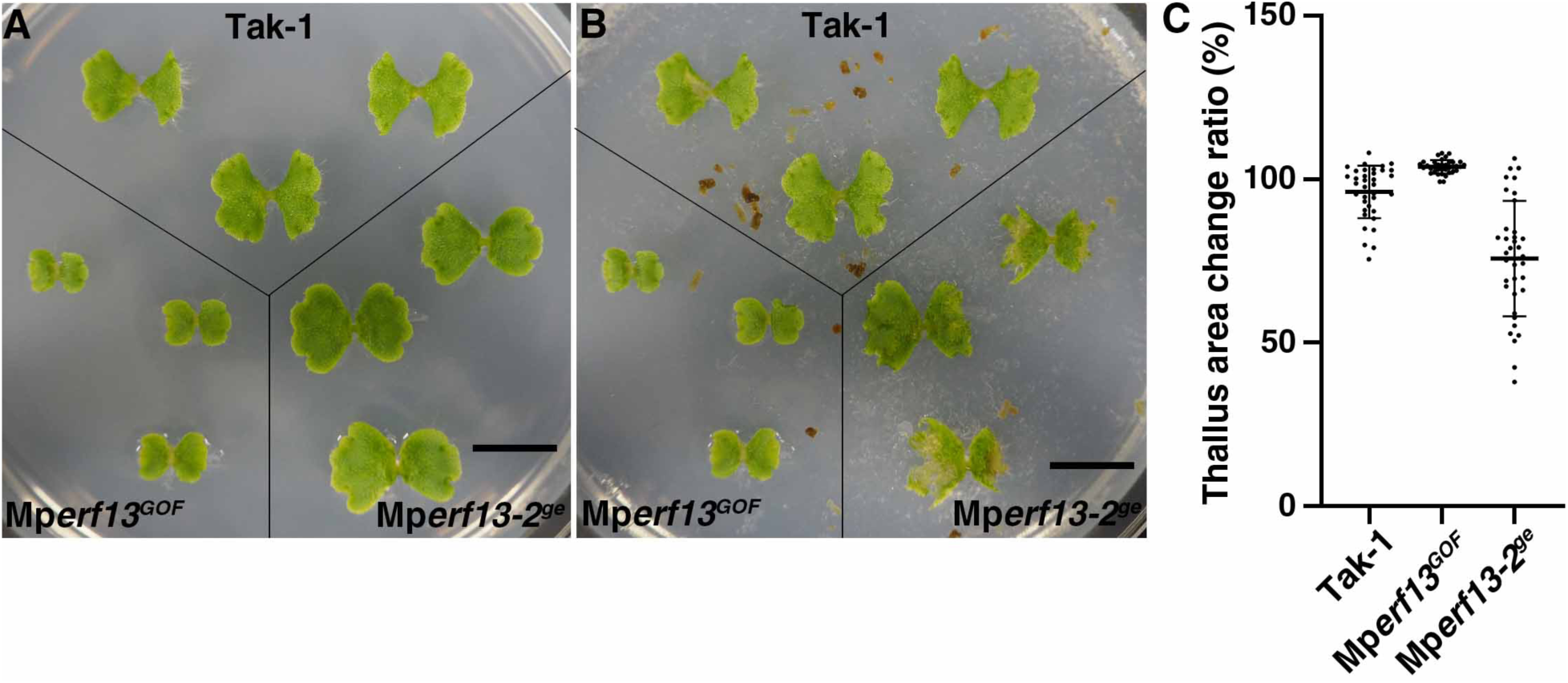
Pill bug feeding assay. (A and B) Ten-day-old thalli of Tak-1, Mp*erf13*^*GOF*^ and Mp*erf13-2*^*ge*^ (A) were co-cultivated with starved pill bugs for 24 hr (B). Bars = 1 cm. (C) Ratios of thallus area before and after co-cultivation with pill bugs were calculated (n = 36 thalli for each genotype). Bars indicate means ± SD. Statistical analyses between Tak-1 and each genotype were conducted with a two-tailed Welch’s *t*-test. The *p*-values were 1.63×10^−5^ for Mp*erf13*^*GOF*^ and 4.91×10^−6^ for Mp*erf13-2*^*ge*^.

**Table S1**. The list of DEGs in Mp*erf13* mutants identified in RNA-Seq analysis.

**Table S2**. List of primers used in this study.

**Movie S1. Formation of clathrin-coated vesicles at the oil body membrane**.

A time-lapse movie of an oil body cell expressing MpCLC1-Citrine. Green and blue pseudo-colors represent fluorescence from Citrine and chlorophyll, respectively. 32× real time. Bar = 5 µm.

**Movie S2. Expression of MpSYP12B in a Marchantia thallus**

A five-day-old thallus expressing 2×YFP driven by the Mp*SYP12B* promoter observed by light sheet microscopy. Green and blue pseudo-colors represent fluorescence from Citrine and chlorophyll, respectively. Bar = 400 µm.

